# Multiple sequential prediction errors during reward processing in the human brain

**DOI:** 10.1101/2020.10.20.347740

**Authors:** Colin W. Hoy, Sheila C. Steiner, Robert T. Knight

## Abstract

Recent developments in reinforcement learning, cognitive control, and systems neuroscience highlight the complimentary roles in learning of valenced reward prediction errors (RPEs) and non-valenced salience prediction errors (PEs) driven by the magnitude of surprise. A core debate in reward learning focuses on whether valenced and non-valenced PEs can be isolated in the human electroencephalogram (EEG). Here, we combine behavioral modeling and single-trial EEG regression revealing a sequence of valenced and non-valenced PEs in an interval timing task dissociating outcome valence, magnitude, and probability. Multiple regression across temporal, spatial, and frequency dimensions revealed a spatio-tempo-spectral cascade from valenced RPE value represented by the feedback related negativity event-related potential (ERP) followed by non-valenced RPE magnitude and outcome probability effects indexed by subsequent P300 and late frontal positivity ERPs. The results show that learning is supported by a sequence of multiple PEs evident in the human EEG.

## INTRODUCTION

Adaptive decision making requires predicting mappings between stimuli, actions, and outcomes to optimize choices that maximize reward. In predictive coding frameworks, these associations are learned and updated via prediction errors (PEs) following surprising feedback. Foundational systems neuroscience findings on reward processing report that transient firing rate modulations of midbrain dopamine (DA) neurons encode reward prediction errors (RPEs), or the valenced difference (either positive or negative) between observed and expected reward values ^1,2^. In parallel, early work in reinforcement learning (RL) established that model-free algorithms such as temporal difference learning can account for basic reward learning phenomena by using RPEs to update the expected value of a stimulus without modeling the influence of actions ^3^.

Recent animal investigations of reward processing and cognitive control emphasize a multitude of learning signals linked to reward and motivation ^4^. Optogenetic dissections of reward circuits report DA responses to both aversive outcomes and salient stimuli regardless of valence ^5–7^, and that separate subpopulations of midbrain DA neurons code for sensory, cognitive, and reward variables {Engelhardt.2019}. Current model-based RL algorithms incorporate non-valenced predictions of states, actions, and outcomes to account for complex behaviors in dynamic environments ^8–11^. Similar predictive coding of non-valenced action-outcome associations is employed by recent models of motor and cognitive control ^12^the predicted response-outcome (PRO) model asserts that the medial prefrontal cortex (MPFC) controls action selection by tracking salient or unexpected outcomes independent of valence ^13^. However, the neural dynamics of these learning signals are poorly understood.

Electroencephalography (EEG) research on reward processing has also identified multiple learning-related event-related potentials (ERPs) ^14^, but their specific relationships to reward and control PEs are unclear. The extant literature focuses primarily on the feedback-related negativity (FRN, also known as the reward positivity; see Proudfit^15^ for details), an early, post-feedback, theta frequency (4-8 Hz) ERP localized to control circuits in MPFC ^16–19^ that has been linked to abnormal dopamine and reward sensitivity in psychiatric conditions ^20,21^. A prominent RL theory proposes the FRN represents a quantitative RPE driven by midbrain DA projections to MPFC ^22^. This hypothesis posits larger, more negative FRN amplitude for more negative RPEs, and also predicts FRN sensitivity to magnitude and probability of rewards, though a recent meta-analysis found mixed evidence for these claims ^23^. Reports of larger FRNs to unexpected positive outcomes ^24–26^ led to the alternative salience theory proposing the FRN represents the degree of surprise of an outcome, similar to non-valenced action-outcome PEs underlying the PRO model. This account predicts the FRN should show a strong effect of non-valenced RPE size driven by unlikely outcome magnitudes or probabilities, but no effect of valence. For example, the salience theory predicts a larger FRN after unlikely than likely wins, whereas the RL theory predicts a smaller FRN after unlikely than likely wins due to more positive RPEs.

One of the main challenges in tackling this core issue in reward learning is dissociating the FRN from overlapping ERPs like the P300, which also co-varies with both valenced and non-valenced features of reward learning ^27–29^. The FRN is often measured by averaging over time using epochs that include the P300 ^14,23^, and early P300 ramping influences the positive peak preceding the FRN ^25^, confounding its use as a reference point in a commonly used peak-to-peak FRN metric. This overlapping component problem is typically addressed by subtracting ERPs across conditions to create difference waves, but isolating individual variables via binary contrast logic does not account for interactions between other components, making it poorly suited to unravel the multiplicity of learning signals and ERPs.

Here we address this controversy by using behavioral modeling of valenced RPEs and non-valenced magnitude and probability signals and applying single-trial EEG regression across temporal, spatial, and frequency dimensions to separate overlapping components. These analyses dissociate valenced RPE value represented by the FRN from non-valenced RPE magnitude indexed by the P300 and also reveal a later frontal positivity tracking outcome probability. This sequence of prediction errors confirms the RL theory of the FRN, and suggests reward processing in the human brain should be reframed as an evolution of multiple learning signals that can be disentangled using single-trial EEG.

## RESULTS

We collected and analyzed EEG from 32 cognitively normal young adults encompassing initial (n = 15) and replication (n = 17) cohorts. Participants performed an interval timing task designed to dissociate the key variables underlying the debate between RL and salience theories: outcome valence, magnitude, and probability. At the beginning of each trial, participants saw a target zone cue whose size indicated the temporal range of responses tolerated as correct (Fig. 1A). Participants then estimated the temporal interval by means of extrapolation from visual motion, and received audiovisual feedback indicating their reaction time (RT) and whether it was within or outside the tolerance (i.e. a win or loss). After each trial, the error tolerance was titrated by two staircase algorithms (Fig. 1B) to clamp accuracy at 82.7 ± 1.7% and 18.1 ± 2.5% (mean ± SD) in easy and hard blocks, respectively (Fig. 1C). This design dissociates outcome valence and probability to separate valenced and non-valenced RPE features by comparing surprising wins and losses. Neutral outcomes with no RT feedback were also delivered on a random subset of 12% of trials to manipulate outcome magnitude as another source of surprise.

**Figure 1:**
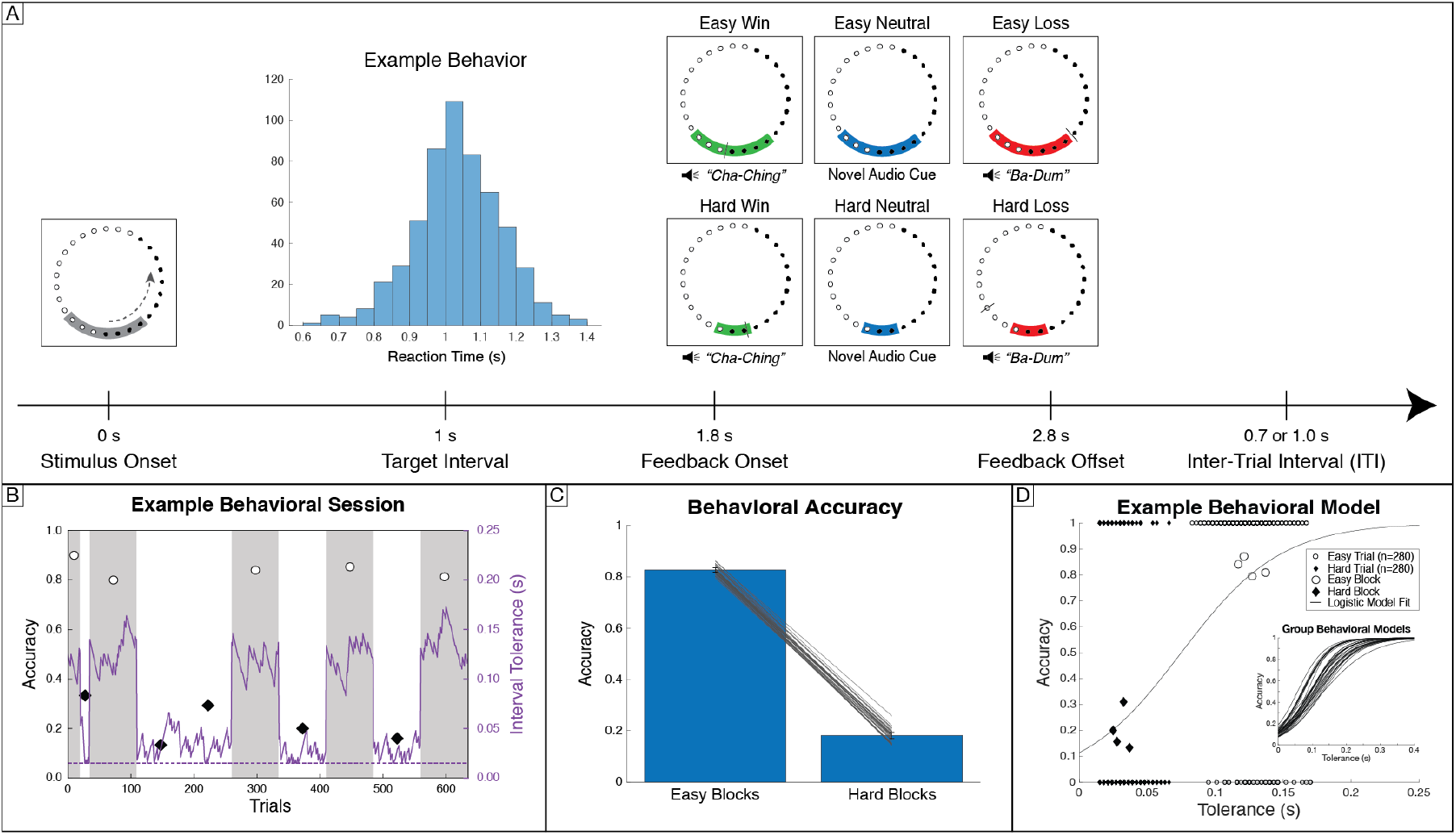
Task design, performance, and behavioral modeling of prediction errors. (A) Participants pressed a button timed to the estimated completion of lights moving around a circle. The gray target zone cue displayed error tolerance around the 1 s target interval. An example participant RT distribution is centered at the target interval. Audiovisual feedback is indicated by the tolerance cue turning green for wins and red for losses. A black tick mark displayed RT feedback. On 12% of randomly selected trials, blue neutral feedback was given with no RT marker. (B) Example recording session for one participant for training (first 35 trials) and experimental blocks. Staircase adjustments of tolerance are plotted in solid purple, and the dotted purple line indicates the minimum bound on tolerance at ±15 ms. Accuracy for easy and hard blocks is plotted as white circles on gray backgrounds and black diamonds on white backgrounds, respectively. (C) Separate staircase procedures resulted in group accuracies of (mean ± SD) 82.7 ± 1.7% for easy and 18.1 ± 2.5% for hard blocks. Error bars indicate standard deviation across participants, with individual participant accuracy overlaid in gray. (D) Tolerance and outcome data for the same example subject. Larger markers show block level accuracy; smaller markers show binary single trial outcomes. Model fit using logistic regression provides single trial estimates of win probability, which can be converted to expected value. Inset shows win probability curves across all participants.

### Behavioral Modeling

To directly compare the predictive power of RL and salience theories, we used computational modeling of individual participant behavior to derive singletrial estimates of valenced RPE value, as well as two sources of salience: non-valenced RPE magnitude and outcome probability. For each subject, we used logistic regression to fit the relationship between the interval tolerance and binary win/loss outcomes across the entire session (Fig. 1D; see inset for group model fits). The resulting model yields the probability of that participant winning for any given tolerance, which was then linearly scaled to the range of rewards (1, 0, and −1 for winning, neutral, and losing outcomes) to quantify expected value for every trial. We then contrast expected value with actual outcomes to obtain single-trial RPE values and derive the absolute value of RPEs to obtain RPE magnitudes. Outcome probability was determined by the frequency of each outcome in each condition. Notably, RPE values for neutral outcomes were non-zero and switched valence across blocks (negative for easy and positive for hard blocks; see model predictions in Supplemental Figure 1A), suggesting they could be interpreted as omissions of the expected outcome.

### Single-trial regression reveals a spatio-temporal cascade from early, frontal RPE value to later, posterior RPE magnitude and frontal probability effects

According to the RL theory, the valenced RPE values derived from our behavioral model should predict FRN amplitude ^22^. In contrast, the salience theory predicts that FRN amplitude should scale with how surprising each outcome is ^13,24^, which increases with non-valenced RPE magnitude and decreases with outcome probability. Importantly, disentangling these three features and resolving the debate between RL and salience accounts depends on addressing the overlapping ERP components problem that confounds traditional point estimates of FRN and P300 amplitude ^27^. Here, we utilize known timing differences between early FRN and later P300 and frontal slow activity by predicting single-trial evoked amplitude at every time point from 50 to 500 ms post-feedback using a multiple regression model combining expected value, RPE value, RPE magnitude, and outcome probability. This analysis was conducted separately at frontal Fz and posterior Pz electrodes to assess the FRN and longer latency positivities.

Grand average ERPs show FRN peaks ~200-250 ms post-feedback at frontal electrode Fz and P300 peaks ~300-350 ms at posterior Pz (Fig. 2A and 2B; see also Supplemental Fig. 6A). The most prominent model result is a large effect of RPE value peaking in the FRN window at 216 ms in electrode Fz (β_max_ = 4.572, *q_FDR_* < 10^-10^; Fig. 3C). In accordance with the RL theory, this positive model coefficient indicates that larger RPE values are associated with smaller amplitude negativities. In other words, larger, more negative FRNs are associated with worse-than-expected outcomes. The RPE value effect decreases as the FRN subsides, and a significant positive RPE magnitude effect emerges. This RPE magnitude effect is maximal in electrode Pz at 308 ms (β_max_ = 1.703, *q_FDR_* < 10^-10^; Fig. 3D), indicating larger non-valenced RPE magnitudes are associated with larger P300 amplitudes.

**Figure 2:**
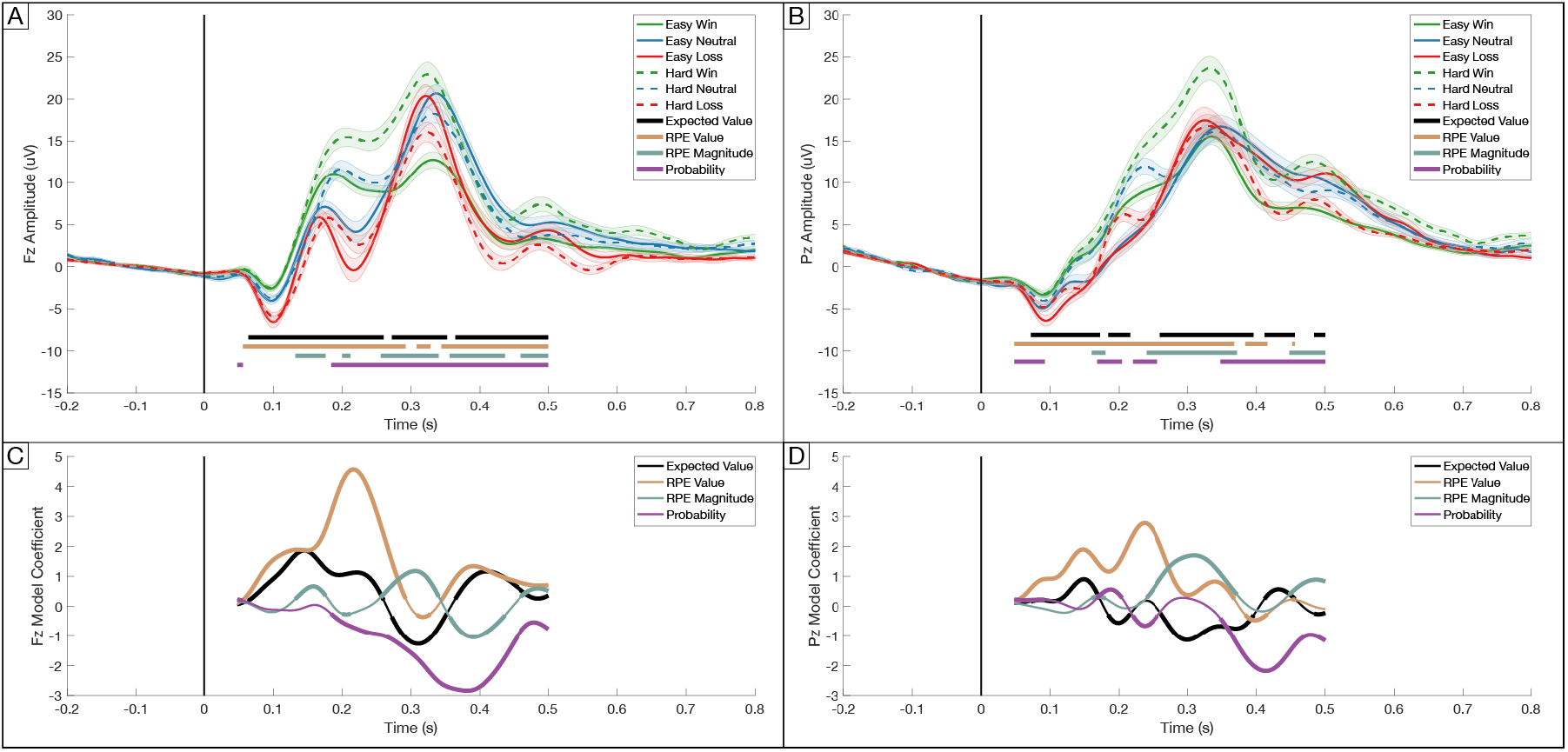
Single-trial modeling of ERP amplitude reveals a sequence of prediction errors. (A) Feedback-locked grand average ERPs at Fz plotted for each condition, with shaded error bars indicating standard error of the mean across participants. Significant periods for each model predictor are marked beneath the waveforms. FRN is evident as prominent negative deflections at Fz ~200 ms post-feedback. (B) Same for parietal electrode Pz, which shows large P300 positivities at ~300 ms. (C) Model coefficients from single-trial multiple regression at each time point from 50-500 ms at Fz show a strong early peak of valenced RPE in the FRN window, followed by later probability effect in the P300 window. Bolding indicates significant time points (*q_FDR_* < 0.05). (D) Same for electrode Pz. Note the increased non-valenced RPE coefficient in the P300 window. See Supplemental Figure 2 for model performance (R^2^), which peaks in the FRN time window for Fz and the P300 time window for Pz.

**Figure 3:**
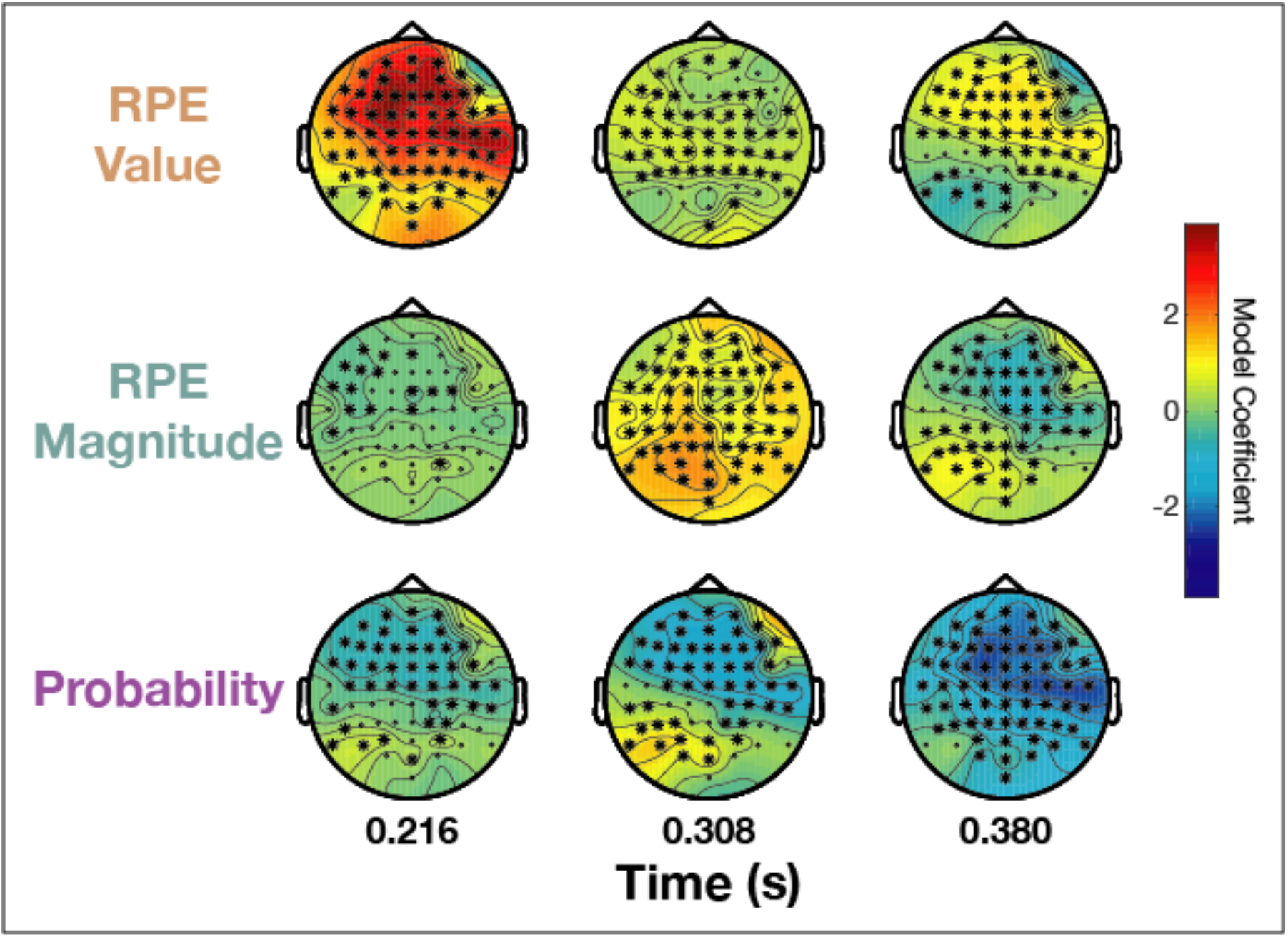
Spatio-temporal dynamics of prediction errors across ERP amplitude scalp topographies. Single-trial regression over all 64 electrodes is computed for three 50 ms windows centered on the largest peak in model coefficients for RPE value (216 ms at Fz), RPE magnitude (308 ms at Pz), and probability (380 ms at Fz) from Fig. 2. Stars indicate significant electrodes (*q_FDR_* < 0.05). Valenced RPE value shows a frontal distribution in the early window (top left). Non-valenced RPE magnitude is maximal at posterior electrodes in the middle window (center). Probability is maximal at fronto-central sensors in the late window (bottom right). See Supplemental Figure 3 for evoked potential topographies.

The temporal coincidence of significant model coefficients for RPE value and magnitude in the epoch between FRN and P300 peaks suggests that previous findings supporting the salience theory could be explained by component overlap confounds, particularly since FRN amplitude is commonly quantified as the mean amplitude from approximately 228-344 ms ^23^, an epoch that encompasses P300 activity. Indeed, Supplemental Figures 1C and 1D show FRN results using traditional mean window and peak-to-peak metrics that confirmed the strong RPE value effect but also show significant probability or RPE magnitude effects, respectively. Importantly, these non-valenced FRN effects using traditional methods were unreliable across replication cohorts.

Lastly, the probability predictor reveals a significant relationship to ERP amplitude that peaks later at 380 ms in Fz (β_max_ = −2.842, *q_FDR_* < 10^-10^; Fig. 3C). Observing this late, frontal positivity in response to unlikely outcomes was possible because few studies have dissociated outcome probability and RPE magnitude as two distinct sources of salience. Collectively, these results suggest a cascade of PEs in reward processing, starting with an early, frontal, valenced RPE value signal in the FRN time window, followed by a later, posterior, non-valenced RPE magnitude effect in the P300 window, and finally a later, fronto-central probability effect.

### Scalp topography dynamics show spatio-temporal segregation of early frontal RPE value, intermediate posterior RPE magnitude, and late fronto-central probability effects

To further disentangle the spatial topographies of sequential PE effects and relate them to known FRN and P300 scalp distributions, we applied our single-trial multiple regression analysis across all electrodes in three 50 ms windows centered on the peaks of each model coefficient from the time-resolved analysis in Fig. 2. Model coefficients for valenced RPE value, non-valenced RPE magnitude, and outcome probability are plotted as scalp topographies in Figure 3 (see Supplemental Figure 3 for evoked amplitude topographies). This analysis confirmed that the largest effect was the valenced RPE value in the early window at anterior frontal sites (β_max_ = 3.887 in 216 ms window at electrode F1, *q_FDR_* < 10^-10^), which then dropped off in magnitude in the middle window before a slight resurgence in the late window. The non-valenced RPE magnitude effect was maximal in the middle 308 ms window at posterior parietal electrodes (β_max_ = 1.859 at electrode PO3, *q_FDR_* < 10^-10^). Finally, the probability effect was focused in fronto-central electrodes in the later 380 ms window (β_max_ = −2.523 at electrode C4, *q_FDR_* < 10^-10^). The spatio-temporal distributions of these effects confirms the association between valenced RPE value and the early FRN epoch in frontal electrodes ^30^, while non-valenced RPE magnitude shows a posterior, parietal distribution matching the P3 0 0 ^28^, and probability appears in fronto-central electrodes long after the FRN time window.

### Single-trial regression of time-frequency power dissociates RPE value in theta frequencies from RPE magnitude and probability effects in delta frequencies

Although the FRN and P300 are defined as ERP phenomena, their waveform characteristics are associated with theta (4-8 Hz) and delta (1-4 Hz) frequencies, respectively ^19,28,31^. To further dissociate contributions of RPE value, RPE magnitude, and probability to these overlapping components, we extracted feedback-locked time-frequency evoked power at Fz (Supplemental Figure 4) and Pz (Supplemental Figure 5). Single-trial multiple regression with our PE model revealed a strong negative relationship between RPE value and theta power (β_max_ = −1.405 at [292 ms, 6 Hz] in electrode Fz, *q_FDR_* < 10^-10^; Fig. 4A). The delayed peak of this theta effect relative to the FRN latency may be due to temporal smoothing inherent in wavelet analyses (500 ms for 3 cycles at 6 Hz) spanning across two RPE value effects in early FRN and later ~400 ms epochs (see theta frequency fluctuations of RPE value coefficients in Fig. 2C). In contrast, RPE magnitude significantly predicted posterior delta power (β_max_ = −0.447 at [260 ms, 3 Hz] in electrode Pz, *q_FDR_* = 5.05 * 10^-10^; Fig. 4B), consistent with the upward ramp of the P300. Probability best predicts 4 Hz power at 392 ms post-feedback (β_max_ = −0.871 in electrode Fz, *q_FDR_* < 10^-10^; Fig. 4A). Overall, more negative RPEs predicted stronger theta power in accordance with the RL theory of the FRN, while delta power associated with the P300 increased with larger RPE magnitudes and more unlikely events.

**Figure 4:**
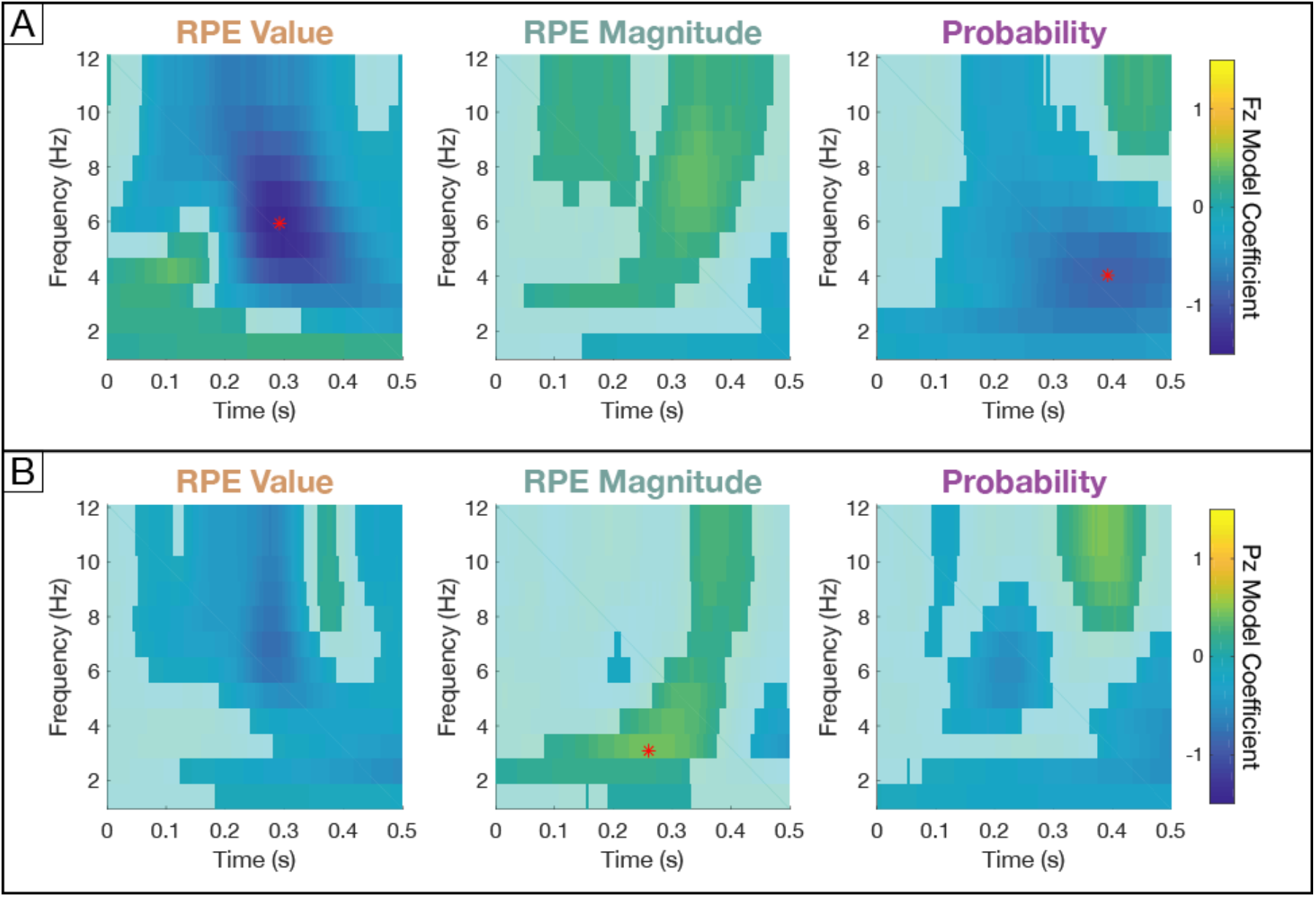
Time-frequency power signatures of prediction errors. (A) Model coefficients fit to evoked power at each time-frequency point for frontal electrode Fz, with non-significant points (q_FDR_ > 0.05) plotted opaquely. Red stars indicate maximal coefficients for each model predictor across both electrodes. Valenced RPE value coefficients peak in frontal theta power, and non-valenced probability coefficients peak later in frontal delta power. (B) Same for Pz. Non-valenced RPE magnitude coefficients peak in posterior delta power.

### FRN peak latency and frontal theta phase are modulated by RPE value

In addition to overlapping components, ERP research must contend with waveshape variability within components due to differences across stimuli, task demands, and individuals ^27,32,33^. For instance, examination of the grand average FRN waveshapes in Supplemental Figure 6A reveals shifts in the latency of the negativity, with losses peaking earlier than wins. Interestingly, neutral outcomes with identical reward values have earlier peak latencies in easy than hard blocks (see inset in Supplemental Figure 6A for direct comparison), despite explicit instructions stating neutral feedback is random and does not reflect performance. Moreover, subjective ratings from post-experiment surveys confirmed participants had neutral overt feelings towards these outcomes in both easy (mean ± SD on zero-centered 9-point Likert scale: −0.2 ± 1.3; *t*(23) = −0.53, *p* = 0.60) and hard (−0.4 ± 1.6; *t*(23) = −1.00, *p* = 0.33) conditions. However, in our behavioral model, neutral outcomes had opposite valence based on contextual expectations, with negative RPE values for omitted wins in easy blocks and positive RPE values for omitted losses in hard blocks.

To determine whether FRN timing shifted systematically according to our PE model predictors, we applied the same multiple regression framework to predict FRN peak latencies. In a general linear model, the only model features predictive of FRN peak latency were RPE value (β = 0.008, *q_FDR_* = 2.80 * 10^-8^; Supp. Fig. 6B) and expected value (β = 0.004, *q_FDR_* = 0.047), which are the two predictors encoding valence. This latency shift in evoked potentials also manifests as biases in theta frequency phase angles and inter-trial phase clustering (ITPC), which measures the consistency of phase angles across trials. Single-trial circular-linear regression between phase angles and individual RPE features localizes this RPE value effect on latencies to theta frequencies in the FRN window (β_max_ = −0.829 at [240 ms, 5 Hz], *q_FDR_* < 10^-10^; Supplemental Fig. 6C). Jack-knife methods were used to estimate the contributions of individual trials to ITPC, and multiple regression revealed more consistent phase angles for positive RPEs at 160 ms in 4 Hz frequencies when early FRN deflections increase phase variability in negative RPE trials (β_max_ = 2.16 * 10^-4^, *q_FDR_* < 10^-10^; Supplemental Fig. 6D). This effect reversed when later FRN deflections in positive RPE trials occurred, such that ITPC was greater for negative RPE trials at 256 ms in theta frequencies (β_max_ = −2.56 * 10^-4^, *q_FDR_* < 10^-10^). This dissociation between subjective ratings and neural signatures of valence suggests our approach can index brain states not revealed by explicit subject report. This finding supports assertions that the FRN is generated by a habitual, model-free reward learning circuit that bypasses conscious representations of goal-directed task structure and instead relies on simpler, implicit associative learning mechanisms ^34^.

## DISCUSSION

These results reveal that early FRN amplitude and peak latency and frontal theta power and phase represent a valenced, quantitative RPE, and that subsequent P300 and posterior delta band activity index non-valenced RPE magnitude. Finally, we observe an even later fronto-central positivity that tracks outcome probability. These observations provide evidence that reward processing signaled in the human EEG is not a unitary event but rather unfolds as a sequence of potentials tracking different elements of PEs.

Our behavioral modeling and regression approach improves on traditional ERP methods in several ways. First, we directly compared theoretical predictions by combining single-trial behavioral estimates of different PEs in mixed-effects multiple regression models enhancing statistical power ^35,36^. Second, single-trial regression at each time(-frequency) point and electrode location provides the high spatio-tempo-spectral resolution needed to disentangle multiple overlapping components ^14,37^. In contrast, traditional mean window and peak-to-peak metrics focus on single measurements influenced by mixtures of components, averaged across trials, and tested at the group level ^23^. For the FRN, these traditional methods replicated the valenced RPE value result, but also found significant non-valenced effects of RPE magnitude or probability depending on the metric, suggesting confounds from overlapping components. Notably, these non-valenced effects using traditional metrics were the only results that did not replicate across two cohorts, potentially due to reduced statistical power relative to our single-trial mixed-effects regression analyses.

Conceptually, recent empirical findings in systems neuroscience ^4,38^ and theoretical advances in RL ^8,39^ emphasize the important role of salience in optimizing choices to maximize reward. This suggests integrating RL and salience perspectives could better account for sequences of valenced and non-valenced PEs as observed in our results ^12,40–42^. Optogenetic dissections of reward circuits have revealed a large diversity of DA signals beyond classic RPEs, including ramping effects ^43^, non-valenced PEs ^44,45^, and salient negative events ^6^. Similarly, single unit recordings in MPFC have identified intermingled populations of single neurons encoding both valenced and non-valenced PEs specific to individual task features (e.g., reward, color, shape) ^46,47^. In parallel, rapid developments in RL have emphasized the role of uncertainty and salience in computing expectations and values to inform action policies ^8^. Collectively, these results suggest that the original RL model of FRN generation via classic DA RPEs projected to MPFC ^22^ needs revision. A realistic model needs to incorporate the diversity of waveshapes, timing, and reward features propagating through mesocorticolimbic pathways that update MPFC control signals driving sequences of valenced and non-valenced effects observable in macroscale EEG recordings.

Indeed, strong links to DA and reward circuits make the FRN a promising biomarker for mood disorders and addiction ^15^,^21^. DA markers predict personality traits like extraversion and sensation seeking ^48^, as well as psychiatric risk (e.g., schizophrenia ^49^, mood disorders ^50^, and addiction ^51^). FRN amplitude also predicts extraversion ^52^, depression symptoms ^53^ and onset ^54^, substance misuse ^55^, and may mediate the relationship between DA and aberrant reward sensitivity in these disorders ^20^. Similarly, P300 abnormalities are observed in a host of disabling neuropsychiatric disorders including substance use disorders ^56^, bipolar disorder ^57^, and schizophrenia ^58^. Refining specific mappings between reward and control PEs and EEG features across multiple dimensions could enhance the power and reliability of these biomarkers and improve their diagnostic and therapeutic potential.

In summary, our experimental design and computational modeling framework support the core prediction of the RL theory that the FRN represents a quantitative valenced RPE ^22^. Multiple regression analyses across temporal, spatial, and frequency dimensions of the data revealed a sequence of subsequent non-valenced PEs in the P300 time window tracking RPE magnitude and outcome probability. These findings demonstrate the power of behavioral modeling and single-trial EEG regression to separate overlapping components, understand the physiology of human behavior, and improve the utility of potential ERP biomarkers for diagnosis and treatment of neuropsychiatric disorders.

## METHODS

### EXPERIMENTAL MODEL AND SUBJECT DETAILS

Data was collected from 41 adult healthy participants (mean ± SD [range]: 20.5 ± 1.4 [18-25] years old; 28 women; 37 right-handed) at the University of California, Berkeley. All subjects reported no history of psychiatric or neurological disorders and had normal, or corrected-to-normal, vision. All subjects were either financially compensated or given course credit and gave written informed consent to experimental protocols approved by the University of California, Berkeley Committees on Human Research.

### METHOD DETAILS

#### Behavioral Task

The Target Time interval timing task was written in PsychoPy ^59^ (v1.85.3) and consisted of eight blocks (four easy and four hard in randomized order) of 75 trials. Following central fixation for an inter-trial interval randomly chosen as 700 or 1000 ms (an earlier task design also included 200 and 400 ms ITIs for four participants in the initial cohort), trials began with presentation of counter-clockwise visual motion from the bottom of a ring of dots at a constant speed to complete the circle at the one-second temporal interval. Participants estimated the interval via button press. The width of a gray target zone indicated the tolerance for successful responses. Veridical win/loss feedback was presented from 1800-2800 ms and composed of (1) the tolerance cue turning green/red, (2) cash register/descending tones auditory cues, and (3) a black tick mark denoting the response time (RT) on the ring.

Participants received ±100 points for wins/losses. Tolerance was bounded at ± 15-400 ms, and separate staircase algorithms for easy and hard blocks adjusted tolerance by −3/+12 and −12/+3 ms following wins/losses, respectively. Participants learned the interval in five initial training trials in which visual motion completed the full circle. For all subsequent trials, dot motion halted after 400 ms to prevent visuo-motor integration, forcing participants to rely on external feedback. Training concluded with 15 easy and 15 hard trials to initialize both staircase algorithms to individual performance levels. Main task blocks introduced neutral outcomes on a random 12% of trials that consisted of blue target zone feedback, a novel oddball auditory stimulus, no RT marker, and no score change.

#### Behavioral Survey

Immediately following the EEG experiment, 24 participants were given a six-question survey to assess their interpretation of example pictures of winning, neutral, or losing outcomes with small or large target zones to indicate easy or hard contexts. In response to the question “How would you feel about this feedback?”, participants rated each outcome on a 9-point Likert scale, where 1 indicated “Terrible!”, 5 indicated “I don’t care...”, and 9 indicated “Great!”. Answers are reported after centering the ratings at the indifference point of 5.

#### Behavioral Modeling

Subjects were excluded because of technical recording errors (n = 4), excessively noisy data (n = 2), or poor behavioral performance (n = 3), leaving 32 participants for analysis. All analyses were piloted on an initial cohort of 15 participants to finalize model parameters and statistical tests before results were replicated in a second cohort of 17 participants. All findings except those in Supplemental Figures 1C and 1D successfully replicated across cohorts, so results presented in the text and figures reflect all 32 datasets combined.

The relationship between the tolerance around the target interval and expected value was fit to individual participant behavior using logistic regression. Specifically, tolerance was used to predict binary win/loss outcomes across trials using the MATLAB function *glmfit* with a binomial distribution and logit linking function. Trials with neutral outcomes were excluded because they were delivered randomly and thus not reflective of performance. The probability of winning (*p_win_)* for each subject was computed as:

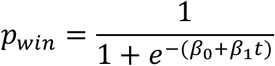

where *β*_0_ is the intercept and *β*_3_ is the slope from the logistic regression, and *t* is the tolerance on a given trial. Expected value was derived by linearly scaling the probability of winning to the reward function ranging from −1 to 1. RPE value was then computed by subtracting expected value from the actual reward value, and RPE magnitude was computed as the absolute value of RPE value. Outcome probability was simply the proportion of each outcome across easy and hard blocks separately.

Notably, this model was fit across all blocks after training under the assumption that subjects learned the task during the 35 training trials, and that the staircase algorithm was appropriately initialized to the subject’s skill level in the training. Since our model is fit using behavior over the entire session, it is possible that it would not describe early trials well, especially if learning occurs over the course of the session. As control analyses, we computed expected value after replacing single-trial win probabilities with block-level accuracy, as well as a rolling average of accuracy on the last 5 or 10 trials. Our single-trial logistic regression model outperformed all of these control models (higher *R*^2^ and lower AIC) for mean window, peak-to-peak, and single-trial amplitude regression analyses.

#### Electrophysiology Recording

EEG data were recorded using a BioSemi ActiveTwo amplifier with a 64-channel active electrode system arranged according to the extended 10-20 system at a sampling rate of 512 Hz. Horizontal electrooculogram (EOG) were recorded from electrodes placed at both outer canthi, and vertical EOG were recorded from an electrode placed below the right eye and right frontopolar electrode FP2. Additionally, two external electrodes were placed on each ear lobe for use in offline re-referencing.

#### Electrophysiology Preprocessing

Preprocessing and analysis used the Fieldtrip toolbox ^60^ and custom code in MATLAB. EEG data were bandpass filtered from 0.1-30 Hz, de-meaned, rereferenced to the average of both ear lobe channels, and then downsampled to 250 Hz. Excessively noisy epochs and channels were removed by visual inspection.

Independent component analysis (ICA) was used to remove artifacts due to channel noise, muscle activity, heartbeat, and EOG (i.e., components correlated with bipolar derivations of horizontal or vertical EOG signals bandpass filtered from 1-15 Hz). Trials were segmented from −0.15 to 2.8 s relative to trial onset, and missing channels were interpolated from neighboring channels via Fieldtrip function *ft_channelrepair*. Final quality checks rejected trials for excessive variance or behavioral outliers (RTs missing, < 0.6 s, or > 1.4 s), resulting in trial counts ranging from 448-524 (mean ± SD: 498.4 ± 20.1).

#### Event-Related Potentials

EEG data were re-aligned to feedback onset and cut to −0.2 to 1 s. ERPs were calculated for each participant by bandpass filtering from 0.5-20 Hz, baseline corrected by subtracting the mean of 200 ms immediately preceding feedback, and averaging across trials.

#### FRN Point Estimates and Latency

To facilitate comparisons with prior FRN studies, we computed traditional mean window and peak-to-peak point estimates of FRN amplitude at electrode Fz. The mean window metric was calculated as the mean amplitude of each participants’ condition-averaged ERPs in a 100 ms window centered on that participant’s FRN peak latency computed across all conditions. The peak-to-peak metric was calculated by subtracting the FRN peak amplitude from the amplitude of the preceding positivity for each condition and subject. To account for variability in ERP waveshapes at the single participant level, peak-to-peak amplitude was only computed if a positive peak was found in the interval 100-260 ms post-feedback that preceded a negative peak in the interval 180-300 ms. According to these criteria, peak-to-peak amplitude could not be computed on 11/192 ERPs. Additionally, the latency of the negative peak in this analysis was used as the FRN peak latency, which was then normalized within participant by subtracting the mean latency across all conditions.

#### Time-Frequency Representations

EEG data were re-aligned and segmented from −0.2 to 1 s around feedback onset. Spectral decompositions were estimated at each time point using Morlet wavelets with 3 cycles of frequencies from 1-12 Hz in linear 1 Hz steps. Task-evoked power was computed as the square of the magnitude of complex Fourier-spectra and baseline corrected by decibel conversion relative to a 200 ms baseline immediately preceding feedback. Phase estimates were derived from the angle of the same complex Fourier-spectra. Inter-trial phase clustering (ITPC), which measures consistency of phase angles across trials for a given time-frequency point ^61^, is computed as

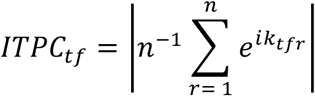

where *n* is the number of trials and *k* is the phase angle at time *t* and frequency *f* on trial *r*. To apply our single-trial regression framework, we used a jackknife procedure to estimate the contribution of each trial to overall ITPC. Specifically, for each trial, ITPC was re-computed excluding that trial and subtracted from overall ITPC computed using all trials.

### QUANTIFICATION AND STATISTICAL ANALYSIS

#### Time-Resolved Modeling

We adopted a multiple linear regression framework to directly compare the predictive power of the valenced and non-valenced PEs derived from our behavioral model. We combined our single-trial model estimates of expected value, valenced RPE value, non-valenced RPE magnitude, and outcome probability in a linear mixed-effects model with random intercepts for each subject to maximize statistical power by accounting for within subject variance. This model was used to predict the temporal evolution of EEG amplitude at each time point from 50 to 500 ms post-feedback using the MATLAB function *fitlme,* which tests significance of model coefficients using two-sided *t*-tests under the null hypothesis the coefficient is equal to zero. Resulting *p* values were corrected for multiple comparisons using false discovery rate ^62^ across time points and model predictors. For clarity, any *p* values corrected for multiple comparisons are reported as *q_FDR_* throughout the manuscript. This analysis was run separately for electrodes Fz and Pz to assess frontal FRN and posterior P300 ERPs.

#### FRN Point Estimates Modeling

This modeling procedure was also used to predict mean window and peak-to-peak FRN metrics measured at electrode Fz. Since these metrics yield one value per condition per participant, each model predictor was averaged within condition for each participant. FDR corrections were applied across model predictors. A nearly identical multiple regression analysis was used to predict FRN peak latency, except the MATLAB function *fitglm* was used without the random intercept for subjects because peak latencies were already normalized within subject.

#### Topography Modeling

To examine the spatial distribution of PE effects on evoked potentials, ERP amplitudes were averaged for all electrodes in three 50 ms windows centered on the largest coefficient from the time-resolved regression for RPE value (216 ms at Fz), RPE magnitude (308 ms at Pz), and outcome probability (380 ms at Fz). The multiple regression model was then used to predict amplitude at each channel in each window, and FDR corrections were applied across all channels, model predictors, and windows.

#### Time-Frequency Power, Phase, and ITPC Modeling

Time-frequency representations were analyzed using the same mixed-effects multiple linear regression model to predict evoked power at each time-frequency point from 0 to 500 ms and 1-12 Hz. FDR multiple comparison corrections were applied across time points, frequencies, and model predictors, again separately for Fz and Pz to assess frontal FRN and posterior P300 effects.

To confirm our FRN peak latency analysis, we used our model to predict phase distributions. Predicting circular variables like phase angles with linear variables like those in our behavioral model requires regressing the linear variable onto both the sine and cosine component of the circular variable, which precludes simultaneous multiple regression using all model predictors. Instead, we predicted single-trial phase angles using separate circular-linear regression analyses for each model predictor with the *CircularRegression* function from the Freely Moving Animal Toolbox (http://fmatoolbox.sourceforge.net/). Since phase values are distributed between 0 and 2π for all participants, no random effect for subject was included in this model. FDR corrections were applied across time-frequency points and regression analyses. Since ITPC is a linear variable, the same procedure as timefrequency power was used to predict single-trial ITPC using our behavioral model.

#### Subjective Ratings

Subjective ratings for neutral trials were tested for significant differences from the indifference point on the 9-point Likert scale after subtracting 5 to center ratings, separately for easy and hard trials. Rating data were tested using two-sided independent samples *t* tests under the null hypothesis that ratings were from a normal distribution with mean equal to zero.

## DATA AVAILABILITY

The datasets generated during and/or analysed during the current study will be made available in the CRCNS (https://crcns.org/) public repository prior to publication.

## SOFTWARE AVAILABILITY

Custom Python and MATLAB code used for preprocessing and analysis is available as a GitHub repository (https://github.com/hoycw/PRlErroreeg), which includes system requirements and dependencies.

## ACKNOWLEDGEMENTS

We thank I. Griffith for help piloting the paradigm, and A. Shah and J. Abbas for help collecting the data. This work was supported by NINDS R37NS21135 and NIMH PO MH109429 (RTK) and NSF GRFP (CWH).

## AUTHOR CONTRIBUTIONS

C.W.H. and R.T.K. designed the experiment. C.W.H. and S.C.S. collected the data. C.W.H. and S.C.S. analyzed the data. C.W.H. and R.T.K. wrote the paper.

## SUPPLEMENTAL FIGURES

**Supplemental Figure 1:**
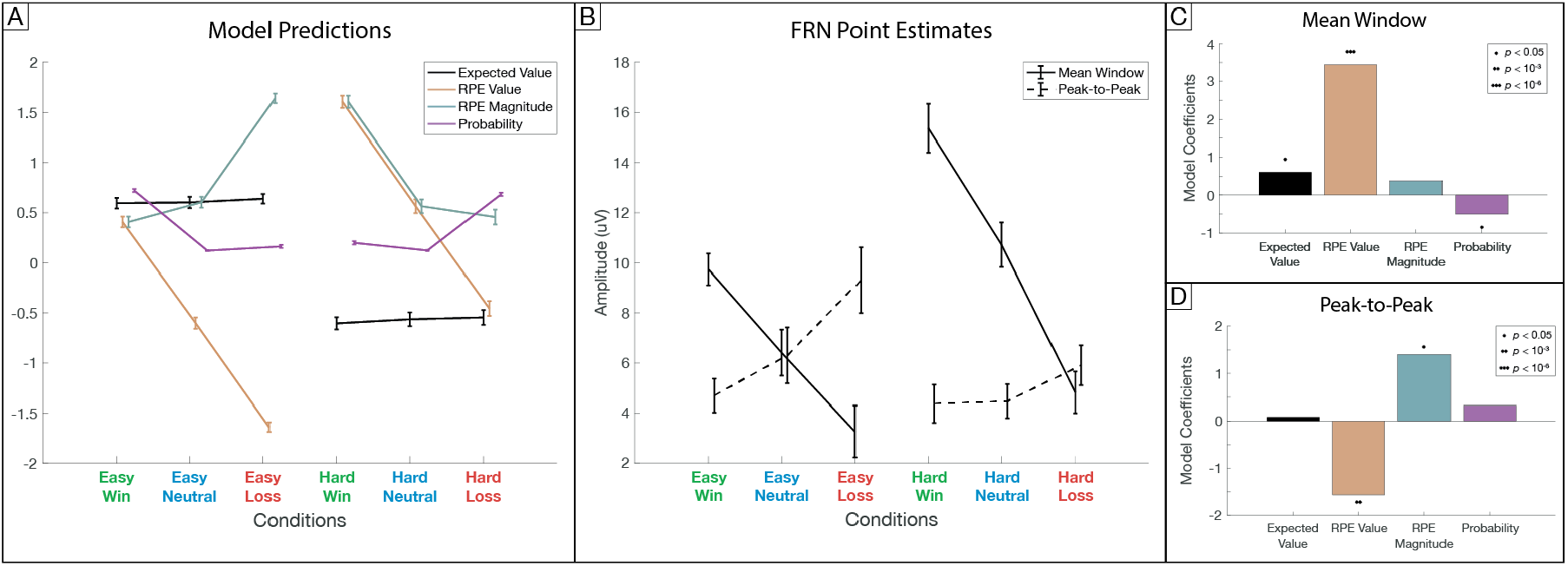
Traditional FRN point estimates track RPE value but also other non-valenced effects. (A) Model predictions by condition. Error bars indicate standard deviation between subjects. (B) Traditional FRN point estimates per condition. Error bars indicate standard error of the mean. Mean window averages single-trial amplitude from 0.178-0.278 s post-feedback (100 ms window centered on the group mean peak latency across all conditions), so larger FRNs have lower mean amplitude. Peak-to-peak calculates differences between amplitudes at FRN and preceding positivity peaks, so larger FRNs have smaller amplitude differentials. (C) Model coefficients from multiple regression of mean window FRN estimates show significance for valenced expected value and RPE value, and also non-valenced probability (q_FDR_ < 0.05). (D) Model coefficients from multiple regression of peak-to-peak FRN estimates are significant (q_FDR_ < 0.05) for valenced RPE value and non-valenced RPE magnitude.

**Supplemental Figure 2:**
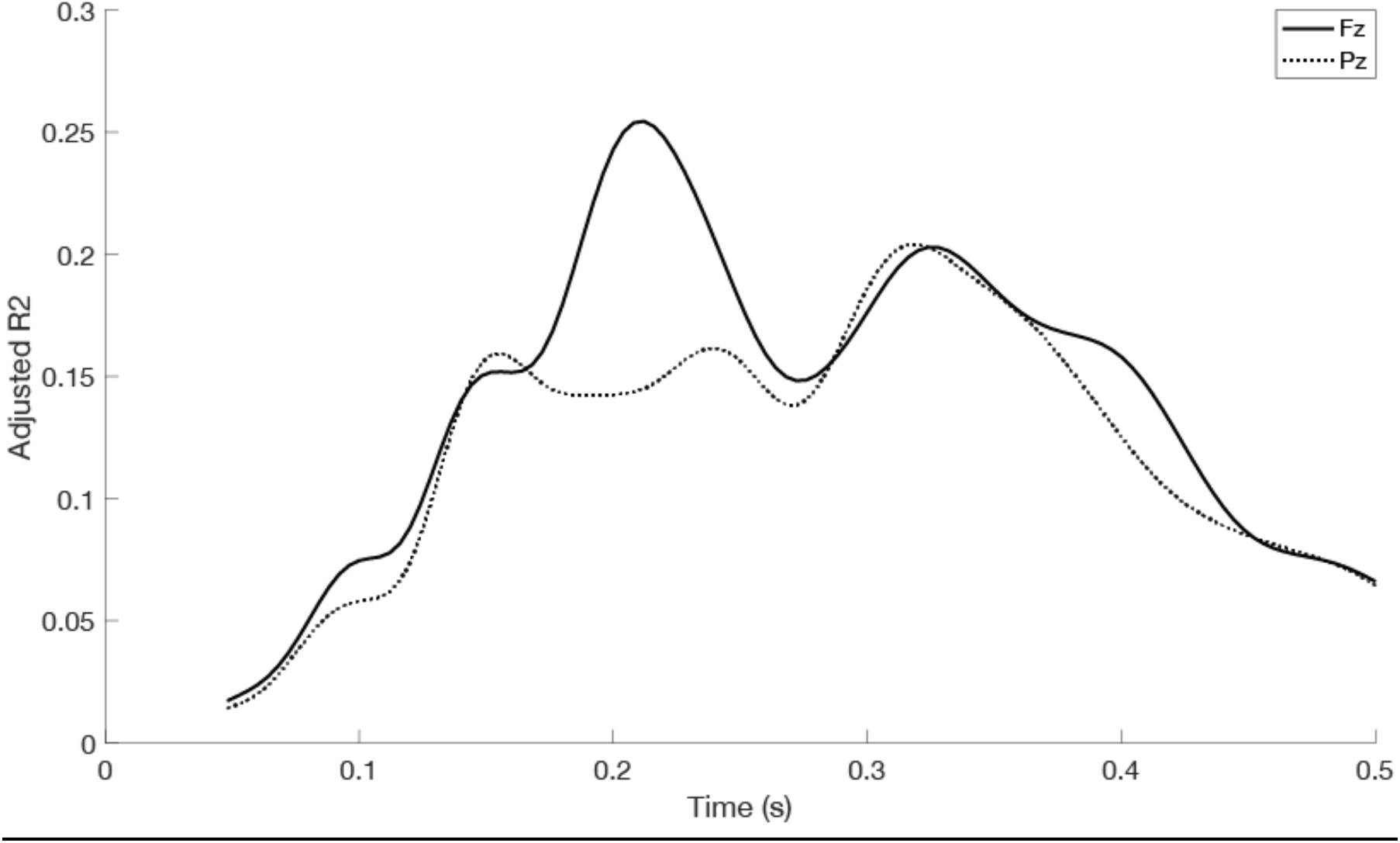
ERP amplitude model fits. (A) Model fits at each time point for frontal electrode Fz plotted as adjusted R^2^. (B) Same for parietal electrode Pz. Note that performance peaks in the FRN time window for Fz (R^2^ = 0.254 at 212 ms) and in the P300 time window for Pz (R^2^ = 0.204 at 320 ms).

**Supplemental Figure 3:**
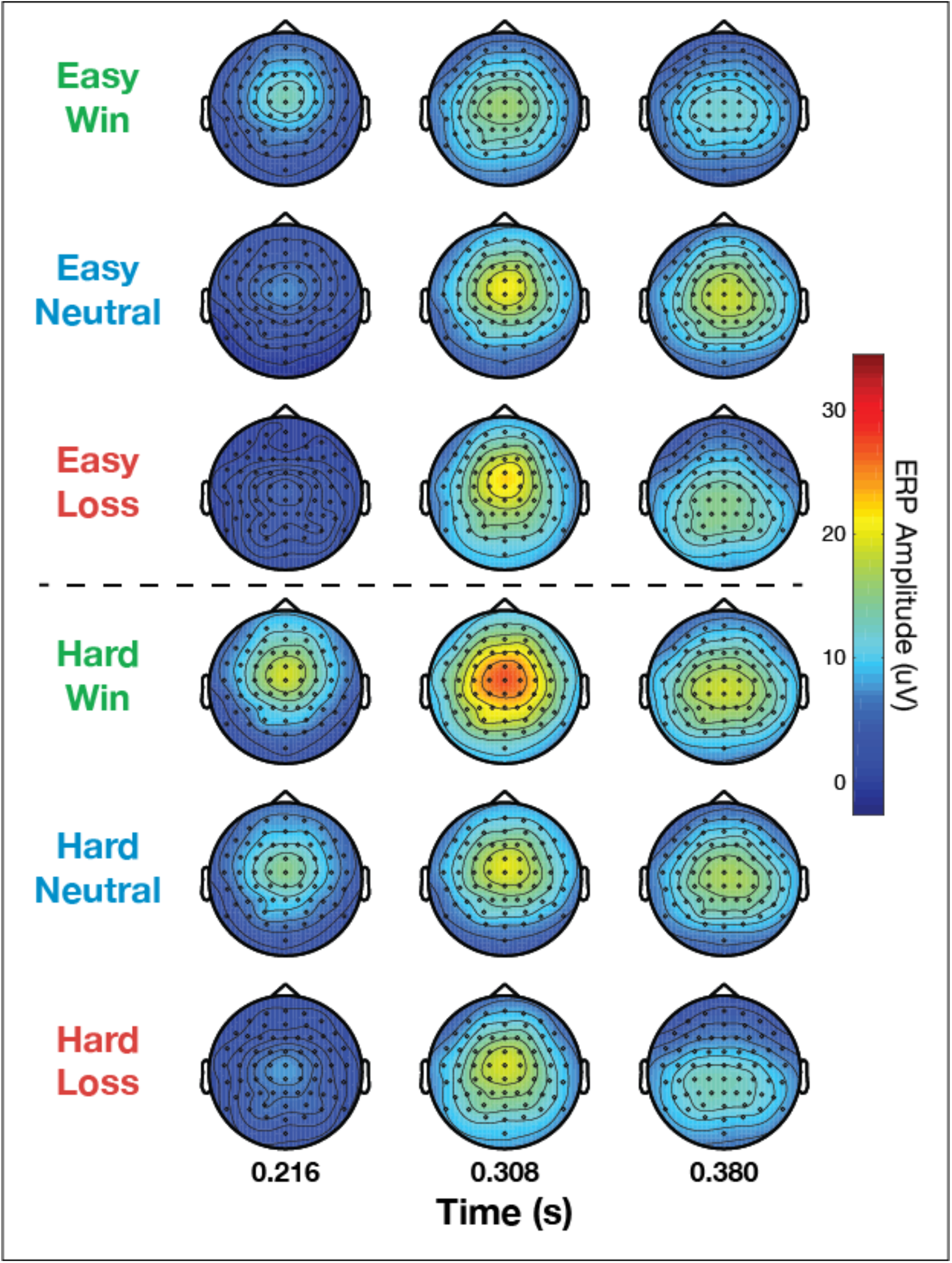
ERP amplitude topography dynamics. ERP amplitudes are averaged within condition and subject in 50 ms windows centered on the peak model coefficients for RPE value (0.216 s), RPE magnitude (0.308 s), and probability (0.380 s) from Fig. 2. Topographies show group averaged amplitude for all electrodes in each window.

**Supplemental Figure 4:**
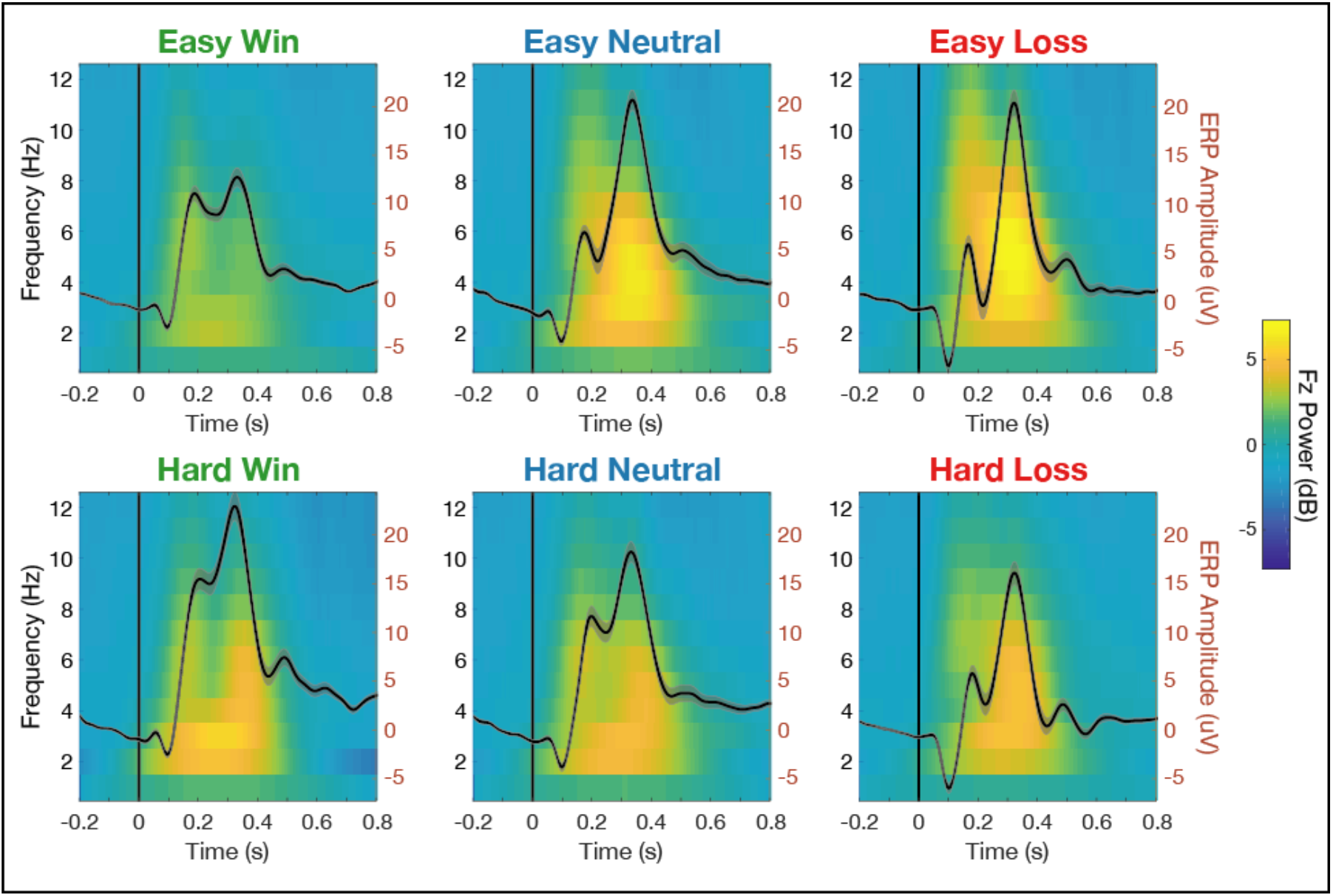
Time-frequency evoked power at Fz for each condition. Grand average ERPs are overlaid on the righty-axis to show the waveshape features driving evoked power.

**Supplemental Figure 5:**
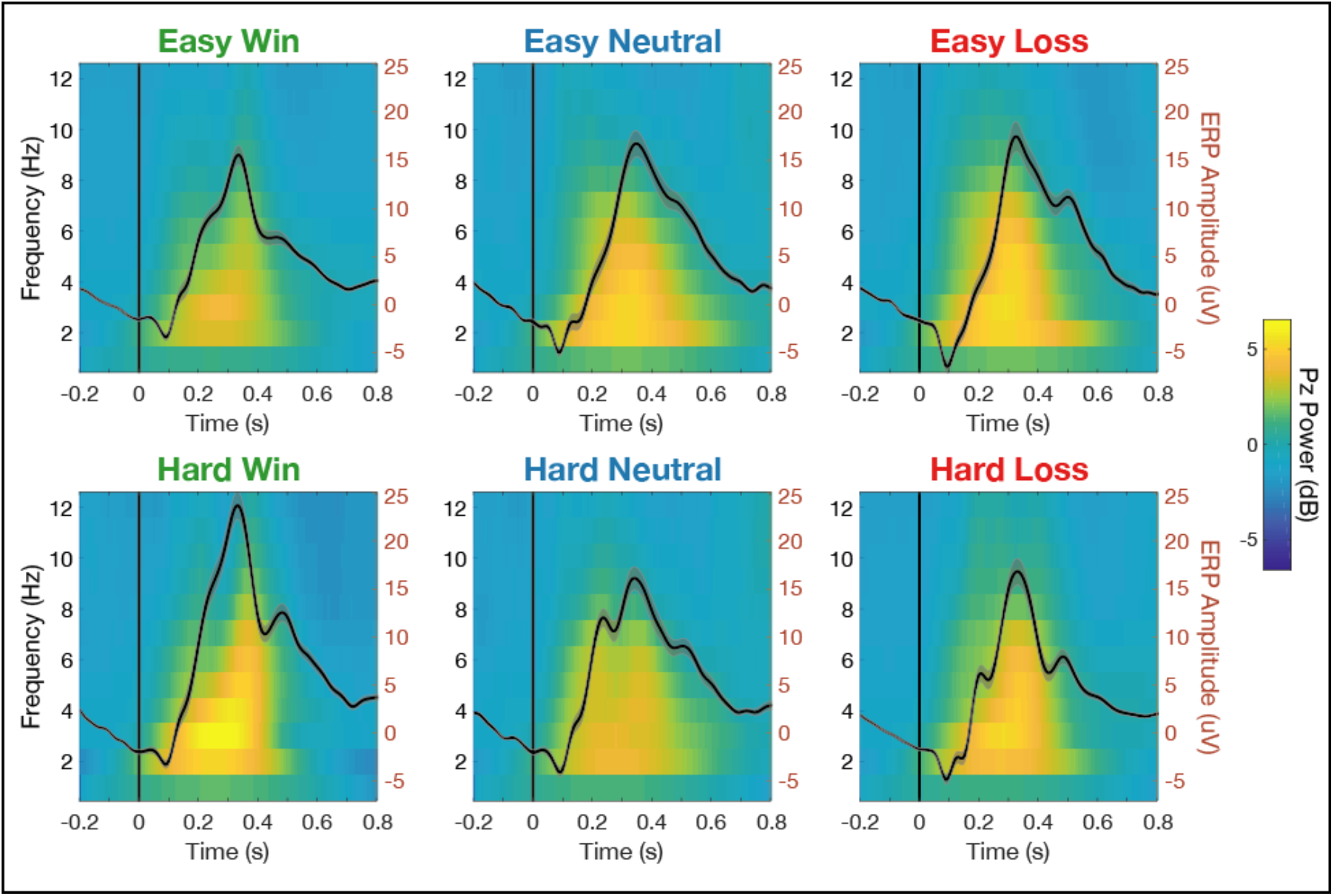
Time-frequency evoked power at Pz for each condition. Grand average ERPs are overlaid on the right y-axis to show the waveshape features driving evoked power.

**Supplemental Figure 6:**
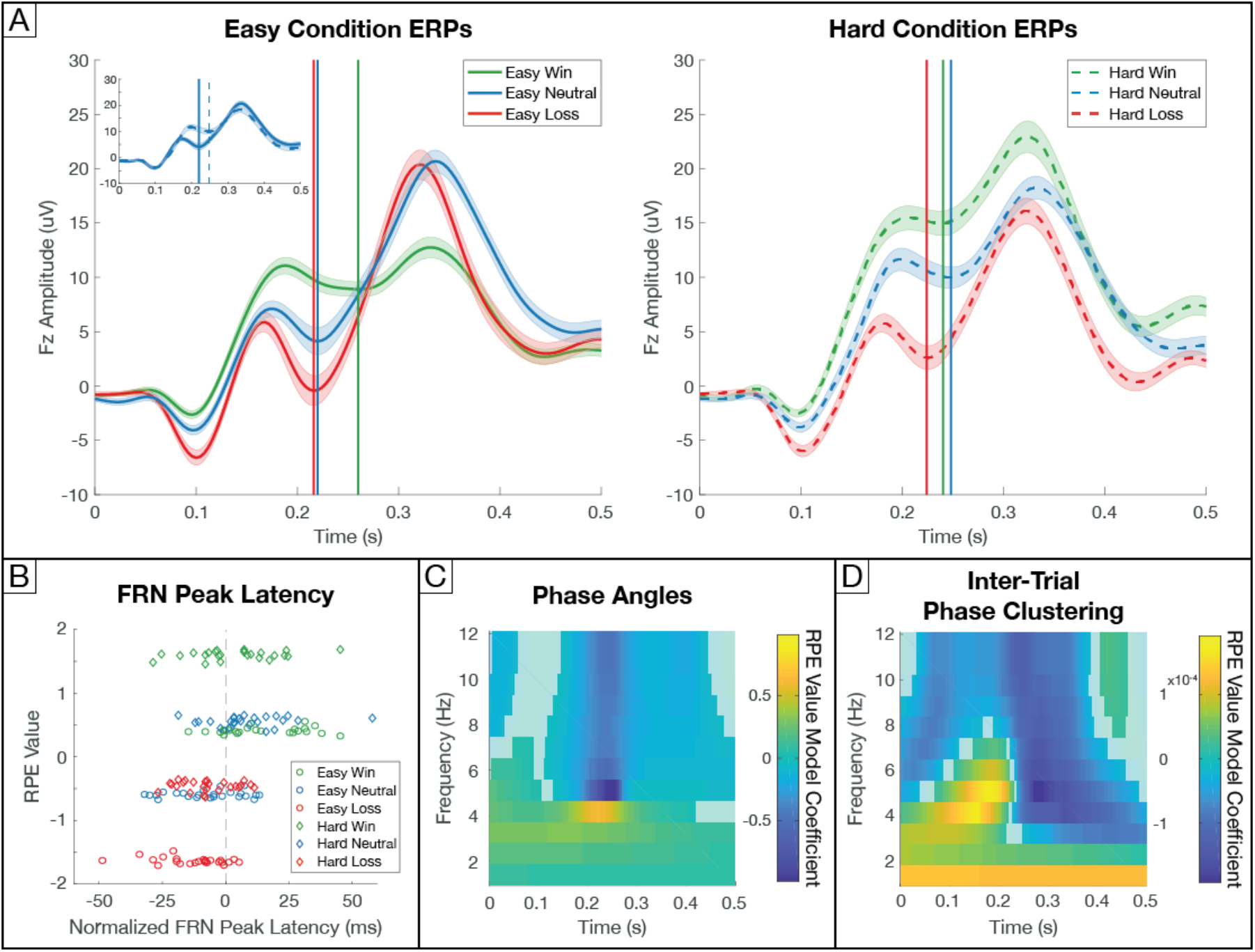
FRN peak latency, phase angles, and phase consistency track valenced RPE value. (A) Grand average ERPs at Fz for easy and hard conditions separately with the FRN peak latency marked by vertical lines. Shaded error bars indicate standard error of the mean across participants. Note that peak latencies for identical neutral outcomes shift early for negative RPEs in easy blocks versus late for positive RPEs in hard blocks (see top left inset for direct comparison). (B) Multiple regression revealed RPE value significantly predicted FRN peak latency (q_FDR_ = 2.80 * 10^-8^). FRN peak latencies for all subjects and conditions are plotted after normalizing for mean FRN latency within subject. Shifts in FRN peak latencies manifest as valenced RPE value encoding in single trial phase (C) and inter-trial phase coherence (D) in the theta frequency range. Opaque points show non-significant time-frequency points (q_FDR_ > 0.05).

